# Kv4.2 regulates baseline synaptic strength by inhibiting R-type channel-mediated calcium signaling in the hippocampus

**DOI:** 10.1101/2023.12.05.570317

**Authors:** Seung Yeon Lee, Minjeong Kwon, Won Kyung Ho, Suk-Ho Lee

**Affiliations:** Cell Physiology Laboratory, Department of Physiology, Seoul National University College of Medicine; Department of Brain and Cognitive Sciences, Seoul National University College of Natural Sciences

## Abstract

Kv4.2 channels, which mediate A-type K^+^ current, exert significant influence on synaptic input signals and synaptic plasticity in the principal cells of the hippocampus. While their influence on activity-dependent regulation of synaptic response is well-established, the impact of Kv4.2 channels on baseline synaptic strength remains elusive. To investigate this, we selectively inhibited postsynaptic Kv4.2 by introducing Kv4.2 antibodies into the hippocampal granule cells and evaluated its impact on the baseline synaptic transmission. Our results demonstrated that Kv4.2 inhibitions led to notable increase in the amplitude of AMPA receptor (AMPAR)-mediated synaptic currents, and this effect was in parallel with the Kv4.2 expression level at dendritic regions. This Kv4.2-dependent synaptic potentiation was effectively abolished by intracellular 10 mM BAPTA or block of R-type calcium channels (RTCC) and downstream signaling molecules including protein kinase A (PKA) and protein kinase C (PKC). Importantly, Kv4.2 inhibitions did not occlude further synaptic strengthening high frequency stimulation, suggesting that synaptic strength regulation by Kv4.2 s distinct from the mechanism of long-term potentiation. Our study highlights the role of Kv4.2 in regulating the baseline synaptic strength, where Kv4.2-mediated inhibition of RTCC is crucial.

## Introduction

The voltage-gated K^+^ channel Kv4.2, responsible for mediating transient A-type currents (I_A_), operates within the subthreshold voltage range to perform various physiological functions. These include the regulation of resting membrane potential (RMP) and action potential (AP) repolarization (Kim et al., 2005) as well as the modulation of synaptic response (Hoffman et al., 1997, Migliore et al., 1999, Johnston et al., 2003, Oule et al., 2021). The graded expression of I_A_ in dendrites acts to constrain the integration and propagation of sub- and suprathreshold voltage signals, including dendritic spikes and back-propagating action potential (b-AP) in the hippocampus (Hoffman et al., 1997, Ramakers and Storm 2002, Johnston et al., 2003, Cai et al., 2004, Losonczy et al., 2006). Moreover, Kv4.2 channels participate in synaptic plasticity by either raising the threshold for LTP induction (Watanabe et al., 2002, Johnston et al., 2003, Kim et al., 2007, Chen et al., 2006, Zhao et al., 2011, Yang et al., 2014, Yang et al., 2015) or undergoing activity-dependent downregulation during long-term potentiation (LTP) (Kim and Hoffman., 2008, Jung and Hoffman., 2009).

It is well established that the regulation of AMPAR densities at postsynaptic density (PSD) is a key mechanism underlying activity-dependent regulation of synaptic strength. Long-term potentiation (LTP) induced by high frequency stimulation requires NMDA receptor activation (Luscher and Malenka., 2012) that facilitate Ca2+ influx to activate Ca2+/calmodulin-dependent kinase II (CaMKII) and various downstream signaling cascades to trigger long-lasting potentiation in synaptic strength (Opazo et al., 2010, Lisman et al., 2012, Sanheuza and Lisman., 2013, Herring and Nicoll., 2016, Yasuda et al., 2022, Hell et al., 2023). However, the regulatory mechanism responsible for maintaining synaptic strength during basal synaptic transmission is not fully understood. Recent findings indicated that genetic deletion and pharmacological blockade of Kv4.2 induce an enhancement in miniature excitatory postsynaptic current (mEPSC) amplitude (Kim et al., 2007, Murphy et al., 2022). These observations suggest the possibility that Kv4.2 plays a role in controlling AMPA receptors (AMPAR) during basal synaptic transmission, although the underlying mechanisms remain to be uncovered. It is therefore crucial to investigate whether Kv4.2 can govern AMPARs to regulate basal synaptic strength, and if so, to elucidate the underlying mechanism.

To address these questions, we introduced Kv4.2-specific antibodies into the cells using whole-cell patch technique and evaluated the role of these channels on basal synaptic transmission at postsynaptic sites. Our findings revealed that Kv4.2 inhibition-induced synaptic potentiation was attributed to an increase in AMPA-mediated current through Ca^+2^-dependent mechanism. Synaptic potentiation by Kv4.2 inhibition required calcium influx through R-type calcium channels (RTCC), and activation of PKC and PKA signaling pathways. Additionally, our observations suggested that Kv4.2 channels may act as input-specific synaptic regulators depending on their expression in the hippocampus. Collectively, we identified Kv4.2 as robust regulator of basal synaptic strength in the hippocampus, potentially contributing to the maintenance of level of AMPA density within physiological range by suppressing RTCC-dependent Ca^2+^ signaling under basal conditions.

## Material and Methods

### Animals

All experiments were performed using C57BL/5 mice of both sex. Animals were housed 5 mice per cage and maintained under specific pathogen-free (SPF) conditions with food and water freely available. The animal maintenance protocols and all experimental procedures were approved by the Institutional Animal Care and Use Committee at Seoul National University (Approval #: SNU-220407-1-7).

### Slice preparation and electrophysiology

Mice were decapitated rapidly after being fully anesthetized by inhalation with isoflurane (Forane; Abott), and the whole brain was immediately removed from the skull, and submerged in iced cold artificial cerebrospinal fluid (aCSF) at 4 °C containing the following (in mM): 110 Choline chloride, 25 NaHCO_3_, 2,5 KCl, 1.25 NaH_2_PO_4_, 0.5 CaCl_2_, 7 MgCl_2_, 10 glucoses, 1 Na-pyruvate, and 0.57 Ascorbate saturating with carbogen (95% O_2_ and 5% CO_2_).

Transverse hippocampal slices (300 μM thick) were prepared using a vibratome (VT1200S, Leica Microsystems). Slices were incubated at 36 °C for 30minutes and the slices were subsequently maintained at room temperature until the recordings in aCSF containing the following (in mM): 125 NaCl, 25 NaHCO_3_, 2.5 KCl, 1.25 NaH_2_PO_4_, 2 CaCl_2_, 1 MgCl_2_, 10 glucoses, consistently bubbled with carbogen (95% O_2_ and 5% CO_2_).

Hippocampal neurons were visualized using an upright microscope equipped with differential interference contrast (DIC) optics (BX51WI, Olympus). Experiments were performed on CA3 pyramidal neurons or mature GCs in the hippocampus. Voltage- and current-clamp recordings were made by the whole-cell patch-clamp technique with EPC-10 USB Double amplitude (HEKA Electronik) at a sampling rate of 10 or 20 kHz. After membrane break-in, two minutes were given to stabilize the neurons.

Patch pipettes (6 – 7 MΩ) and monopolar stimulator pipettes (3 – 4 MΩ) were obtained from borosilicate glass capillaries with a horizontal pipette puller (P-97, Sutter Instruments). The internal pipette solution contained the following (in mM): 143 K-gluconate, 7 KCl, 15 HEPES, 4 MgATP, 0.3 NaGTP, 4 Na-ascorbate, and 0.1 EGTA, with the pH adjusted to 7.3 with KOH, and with osmolality of approximately 300 mOsml/L. To measure IPSP and IPSC amplitude, internal pipette solution contained the Cs-based internal solution in following (in mM): 120 Cs-methanesulfonate, 10 CsCl, 10 HEPES, 2 MgCl, 3 MgATP, 0.4 NaGTP, 5 Na_2_+ phosphocreatine, 0.1 EGTA, with pH adjusted to 7.3 with CsOH. The antibody blocking experiments included Kv4.2 antibodies (1 μg/ml) in the pipette solution. Series resistance (R_s_) after establishing whole-cell configuration was between 10 and 20 MΩ. R_s_ was monitored by applying a short (10s) hyperpolarization (1mV) pulse during the recording. If R_s_ of the recorded cells changed 20% of the baseline value, the cells were discarded.

For voltage-clamp experiments, all recordings were performed at holding potential −70 mV. To record outward K^+^ currents, TTX (0.5 μM), CdCl_2_ (300 μM), and NiCl_2_ (500 μM) were additionally applied to block Na^+^ and Ca^2+^ channels, respectively. Outward K^+^ currents were evoked by injecting +30mV pulse for 1 seconds durations.

For EPSC recordings, the stimulator intensity (100 μs duration; 8-18 V) of extracellular stimulation was adjusted to evoke EPSC amplitudes between 50 pA and 150 pA for the baseline with 50ms intervals delivered every 10s or 30s between sweeps. The holding potential was maintained at −70mV. For EPSP recordings, the cells were held at its RMP. A stimulator (Stimulus Isolator A360; WPI) connected to a monopolar electrode filled with recording aCSF was placed in outer molecular layer of the DG to evoke LEC stimulation induced synaptic responses in the mature granule cells of the hippocampus. Synaptic responses at other synapses are indicated. The PPR was calculated as the ratio of the second EPSC over the first EPSC amplitude. All recordings were performed in the presence of GABA blockers (100 μM PTX, 1 μM CGP52432) to block inhibitory synaptic transmission unless otherwise indicated.

The LTP was induced by applying 10 bouts of high frequency stimulation (HFS) of perforant path synapses. HFS consists of 10 stimuli at 100 Hz under current clamp mode and the bouts were given at 5 Hz. The stimulation intensity was adjusted to evoked at least 2 AP at the first bout.

Intrinsic properties under current-clamp mode and the following parameters were obtained: (1) resting membrane potential (RMP), (2) input resistance (membrane potential changes at a given hyperpolarizing current input (−35 pA, 500 ms)), (3) *F–I* curve (firing frequencies (F) against the amplitude of injected currents from 50pA to 400pA for 500ms duration with 50pA increment (I)), (4) AP half-width (measured as the width at 50% of the spike peak amplitude) (5) AP threshold (the voltage at which d*V*/d*t* exceeded 40 mV/ms).

### Immunohistochemistry analysis

For the immunofluorescence staining, 7-week old C57BL/6 mice were anesthetized with isoflurane and perfused transcardially with 1X PBS and 4% paraformaldehyde (PFA) for 10-15minutes. Brains were removed and cut into 30 μm-thick sections using a vibratome (VT1200S, Leica), and postfixed overnight at 4 °C in 4% PFA solution. The slices were washed 5 times in 1X PBS with 0.3% Triton X-100 (PBS-T) for 5 minutes. Then the sections were incubated three times in a blocking solution (2.5% donkey serum + 2.5% goat serum in PBS-T) for 1 hour at room temperature. The sections were then incubated overnight at 4 °C in blocking solutions containing primary antibodies (anti-Kv4.2 antibody, APC-023, Alomone). After washing 5 times in 0.3% PBS-T for 5 minutes, sections were then incubated with secondary antibodies in blocking solution for at least 1 hr at RT. After rinsing with PBS, sections were mounted on glass slides using mounting medium containing DAPI. The immunostained sections were imaged with a confocal laser scanning microscope (FV1200, Olympus) using a 40x oil-immersion objective lens.

### Data analysis

All data were presented as mean ± standard error of the mean (SEM). Statistical analysis was performed using IgorPro (Version 7.08, Wavemetrics) and Prism. Nonparametric Mann-Whitney U test was used to compare non-paired groups, and Wilcoxon signed-rank test or paired sample t test were used to compare paired groups. Fluorescence intensity was analyzed using Image J. P-values of <0.05 were considered statistically significant.

## Results

### Kv4.2 channel contributes to regulating the synaptic transmission of dentate granule cells

To investigate the functional role of Kv4.2 in regulating synaptic transmission in the dentate gyrus (DG), we performed whole-cell recordings to record synaptic responses in mature granule cells (GCs) in the DG by stimulating lateral perforant pathway (LPP). Throughout the studies, inhibitory synaptic transmission was blocked by PTX (100 μM, GABA_A_R blocker) and CGP52432 (1 μM, GABA_b_R blocker) unless indicated otherwise. Mature GC, characterized by their location within the outer granule cell layer and their low input resistance (R_in_ < 200 MΩ), were selected for observations to avoid potential confounding factors related to the maturation stages of GCs (Kerloch et al., 2019).

To selectively inhibit Kv4.2 channel, Kv4.2 antibodies (Kv4.2 Ab) were introduced via an intracellular pipette solution (1 μg/ml) in GCs, following our previously established protocol (Kim et al, 2020). The specificity of the Kv4.2 Ab in blocking Kv4.2-mediated currents was confirmed by monitoring changes in outward currents evoked by a depolarizing voltage step, as depicted in Figure 1A. Kv4.2 dialysis led to a gradual reduction in the amplitude of transient outward current (Figure 1A, 0.71 ± 0.04, n = 6, p < 0.01), while sustained outward current remained intact (Figure 1A, 1.07 ± 0.04, n = 6, p = 0.11), confirming selective reduction of Kv4.2-mediated transient outward currents (Rhodes et al., 2004).

**Figure 1.**
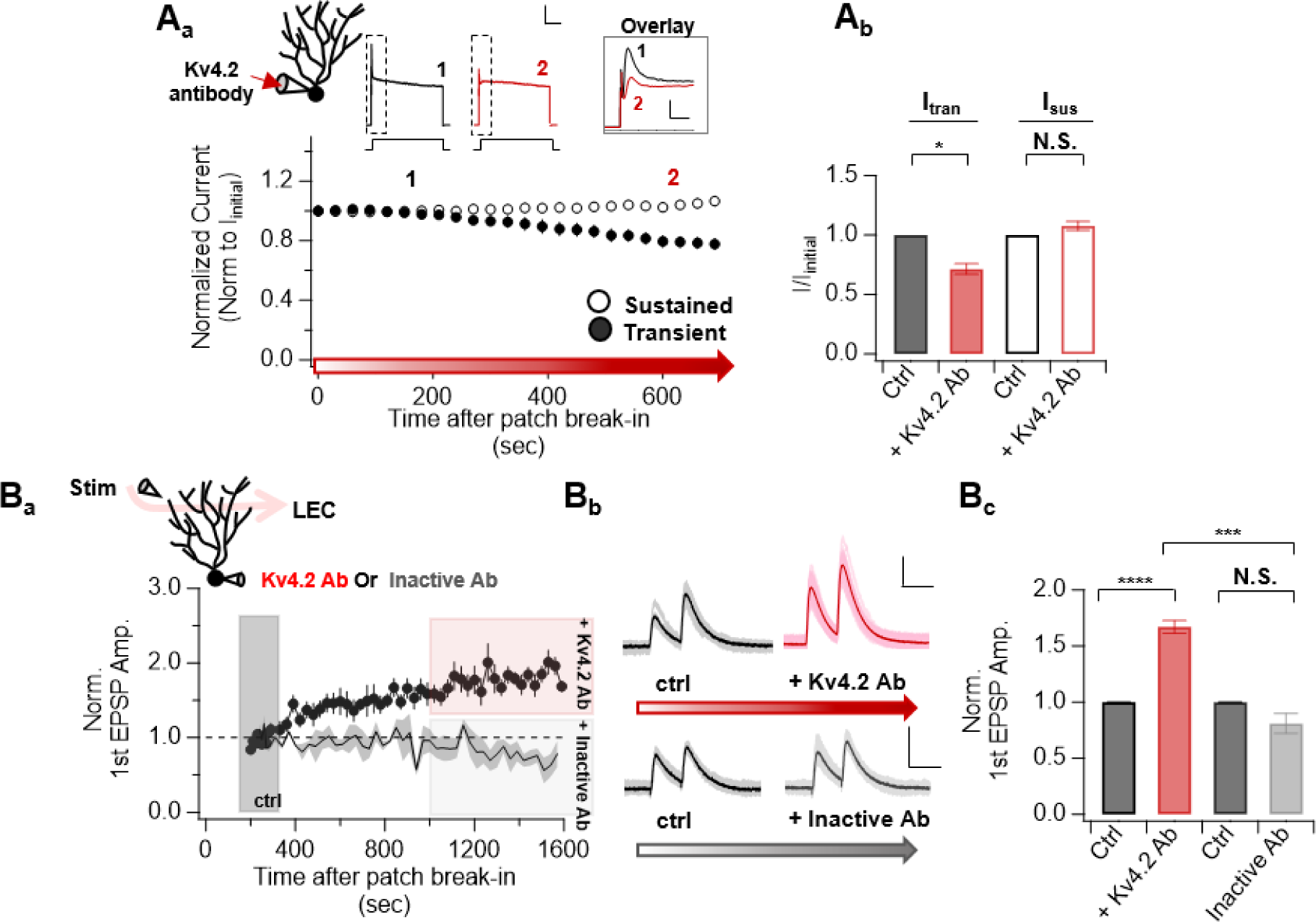
Kv4.2-mediated A-type K+ current enhances EPSP amplitude in mature granule cells. **A_a_** (top) Left: Illustration of the recording configuration, demonstrating the dialysis of Kv4.2 antibodies (Kv4.2 Ab) through the recording pipette in granule cells of the dentate gyrus. Right: Representative traces of outward currents evoked by a depolarizing voltage step pulse from a holding potential of −70mV to + 30mV recorded at 3 min (marked with 1, black) and 15-min (marked with 2, red) after patch break-in from dentate granule cells of the hippocampus. Scale bar, 1nA and 200ms. Inset showing superimposed current traces of 1 and 2 highlighting the specific reduction of transient A-type current due to the dialysis of Kv4.2 antibodies through intracellular solution. Scale bar, 1nA and 10ms. (bottom) The current amplitude, both transient (closed circle) or sustained (opened circle) current, were normalized to their respective initial value following patch break-in. These normalized amplitudes were plotted against the duration of the experiment **A_b_**. I_trans_ or I_sus_ obtained at 15 min after patch break-in were normalized to their respective initial current amplitude and were plotted as bars with SEM **B_a_**. Top: Recording configuration depicting the dialysis of either Kv4.2 Ab or heat-inactivated Kv4.2 Ab through recording pipette, with the stimulating electrode placed in OML region of the dentate gyrus. Bottom: Time course of normalized EPSP amplitude in internal solution with Kv4.2 Ab (represented by closed circles) or heat-inactivated Kv4.2 Ab (represented by lines). EPSP amplitudes were normalized to mean amplitude of the synaptic responses recorded at 3-5 minutes after patch break-in (shaded in black). Kv4.2 Ab induced increased EPSP amplitude while EPSP amplitude remained constant with Inactive Ab. Data are binned into 30ms intervals, mean ± SEM. **B_b_**. Representative average EPSP traces taken the indicated shaded time points during the recording period. Scale bars, 10mV and 50ms. Top: Representative traces of eEPSP amplitude in control (ctrl, black) and under the influence of Kv4.2 Ab effects (+ Kv4.2 Ab, red). Bottom: Representative traces of eEPSP amplitude in control (ctrl, black) and with Inactive Ab effect (+ Inactive Ab, gray). **B_c_**. Bar graph illustrating the enhancement of EPSP amplitude resulting from the inclusion of Kv4.2 Ab in the intracellular pipette, with no observable change when using Inactive Ab All data are represented as mean ± SEM. Statistical significance was evaluated by paired t tests or two-sample t-test. **p < 0.01, ****p<0.000, n.s.=not significant

Subsequently, we examined the effect of Kv4.2 inhibitions on synaptic potentials in LPP-GC synapses (Figure 1B). Notably, intracellular dialysis of Kv4.2 Ab gradually increased the EPSP amplitude to a significant extent, reaching a maximal effect within 15 minutes in most cells (Figure 1B, 1.69 ± 0.05, n = 85, p < 0.0001). Average traces of synaptic responses were obtained within 3-5 minutes (control) and 15-30 minutes (+ Kv4.2 Ab) to provide a comparative view of how Kv4.2 dialysis induced enhancement in EPSP amplitude over time (Figure 1Bb). In contrast, the introduction of heat-inactivated Kv4.2 Ab (Inactive Ab, 1 μg/ml) via intracellular solution did not induce a significant change in EPSP amplitude (Figure 1B, 0.81 ± 0.09, n = 6, p = 0.95), highlighting the role of Kv4.2-mediated I_A_ current in shaping the amplitudes of synaptic response in LPP-GC synapses.

### Kv4.2 channel acts as synaptic strength regulator in dentate granule cells of the hippocampus

We then examined the effect of Kv4.2 Ab on EPSC amplitude in LPP-GC synapses. Remarkably, Kv4.2 Ab enhanced EPSC amplitude by approximately 1.6-fold by Kv4.2 Ab, contrasting with the effect of the Inactive Ab (Figure 2A, EPSC_kv4.2 Ab_: 1.61 ± 0.06, n = 85, p < 0.0001; EPSC_inactive Ab_: 0.99 ± 0.05, n = 6, p = 0.90). This increase in EPSC amplitude was accompanied by a slight but significant decrease in paired pulse ration (PPR) (Figure 2B, PPR_Ctrl_ vs PPR_Kv4.2_: 1.74 ± 0.04 vs 1.49 ± 0.033, n = 5, p < 0.0001). Furthermore, Kv4.2 had no effect on the EPSC amplitude in cells recorded with cesium-based internal solution, indicating the requirement for postsynaptic K^+^ current to induce EPSC enhancement (Figure 2C, 0.98 ± 0.01, n = 5 p = 0.87). Moreover, there was no significant alteration in NMDA-EPSC amplitude in LPP-GC synapses (Figure 2D, 1.13 ± 0.11, n = 8, p = 0.38) or in inhibitory synaptic transmission following Kv4.2 inhibitions (Figure 2E, 2F, IPSC: 1.04 ± 0.06, n = 9, p = 0.49; IPSP: 0.95 ± 0.045, n = 9, p =0.26). These findings underscored the significant contribution of postsynaptic Kv4.2 channels in regulating AMPA-mediated synaptic currents in GCs.

**Figure 2.**
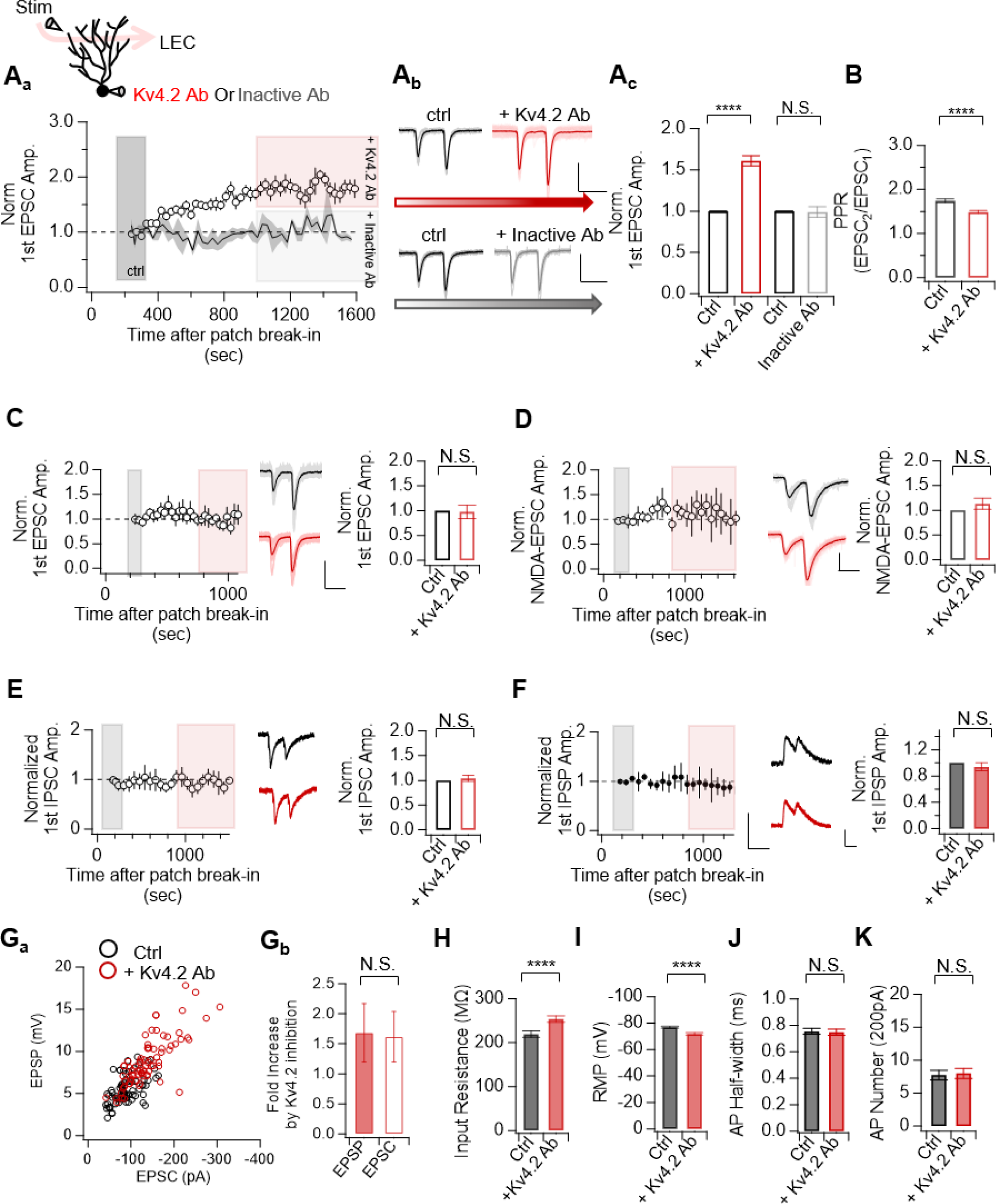
AMPA-dependent synaptic strength is regulated by Kv4.2 channels in dentate granule cells. **A_a_.** Top: Recording configuration depicting the dialysis of either Kv4.2 Ab or heat-inactivated Kv4.2 Ab through recording pipette, with the stimulating electrode placed in OML region of the dentate gyrus. Bottom: Time course of normalized EPSC amplitude in internal solution with Kv4.2 Ab (represented by open circles) or heat-inactivated Kv4.2 Ab (represented by lines). EPSC amplitudes were normalized to mean amplitude of the synaptic responses recorded at 3-5 minutes after patch break-in (shaded in black). Kv4.2 Ab induced increased EPSC amplitude while EPSC amplitude remained constant with Inactive Ab. Data are binned into 30ms intervals, mean ± SEM. **A_b_**. Representative average EPSC traces taken the indicated shaded time points during the recording period. Scale bars, 100pA and 50ms. Top: Representative traces of EPSC amplitude in control (ctrl, black) and under the influence of Kv4.2 Ab effects (+ Kv4.2 Ab, red). Bottom: Representative traces of EPSC amplitude in control (ctrl, black) and with Inactive Ab effect (+ Inactive Ab, gray). **A_c_**. Bar graph illustrating the enhancement of EPSC amplitude resulting from the inclusion of Kv4.2 Ab in the intracellular pipette, with no observable change when using Inactive Ab. **B**. Bar graph illustrating the reduction in PPR by inclusion of Kv4.2 Ab in the intracellular pipette. **C.** (Left) Time course of normalized EPSC amplitude in cesium-based internal solution with Kv4.2 Ab. EPSC amplitudes were normalized to mean amplitude of the synaptic responses recorded at 3-5 minutes after patch break-in (shaded in black). Data are binned into 30ms intervals, mean ± SEM. (Middle) Representative average EPSC traces taken the indicated shaded time points during the recording period. Scale bars, 50pA and 50ms. Representative traces of EPSC amplitude in control (ctrl, black) and under the influence of Kv4.2 Ab effects (+ Kv4.2 Ab, red). (Right) Bar graph illustrating no observable change in EPSC amplitude due to Kv4.2 Ab **D.** (Left) Time course of normalized NMDA-EPSC amplitude in internal solution with Kv4.2 Ab. EPSC amplitudes were normalized to mean amplitude of the synaptic responses recorded at 3-5 minutes after patch break-in (shaded in black). The recordings were conducted in magnesium free solution and in presence of 20μM CNQX, 100μM PTX, 1μM CGP to block AMPA and GABA receptors. Data are binned into 30ms intervals, mean ± SEM. (Middle) Representative average EPSC traces taken the indicated shaded time points during the recording period. Scale bars, 50pA and 50ms. (Right) Bar graph illustrating no observable change in NMDA-EPSC amplitude due to Kv4.2 Ab **E-F.** (Left) Time course of normalized 1^st^ evoked IPSC amplitude (D) or 1^st^ IPSP amplitude (E) using high chloride internal solution containing Kv4.2 Ab. The recordings were conducted in the presence of 20μM CNQX and 100μM PTX to block excitatory synaptic transmission. Data are binned into 30ms intervals, mean ± SEM. (Middle) Representative average IPSC traces taken the indicated shaded time points during the recording period. Scale bars, 50pA and 50ms. (Right) Bar graph illustrating that Kv4.2 inhibition have no influence on IPSC or IPSP amplitude. **G_a._** Ratio of EPSC change (EPSC_Kv4.2_/EPSC_ctrl_) is plotted against EPSP change (EPSP_Kv4.2_/EPSP_ctrl_). Ratio of individual cells are shown in gray and the average ratio is marked in red circle. **G_b_**. Bar graph illustrating equivalent increase in both EPSP amplitude and EPSC amplitude by dialysis of Kv4.2 Ab **H-K**. Intrinsic properties were measured in dentate GCs at 3minutes and at 20minutes after patch break-in to measure Kv4.2 inhibition changes in the following parameters: (F) Input resistance by injecting −30pA current (G) Resting membrane potential (RMP) (H) AP half-width obtained at rheobase current injection for 500ms and (I) number of AP obtained at 200pA current injection All data are represented as mean ± SEM. Statistical significance was evaluated by paired t tests, two-sample t-test. ****p<0.0001, n.s= not significant

It is generally thought that K^+^ channel inhibition can amplify EPSPs by increasing input resistance or dendritic excitability. To examine whether Kv4.2 inhibitions has amplifying effects on EPSPs, we compared the magnitude of increase induced by Kv4.2 Ab on EPSP and EPSC amplitude (Fig. 2G). The results showed equivalent enhancements, indicating that the inhibition of Kv4.2 channels primarily enhanced synaptic strength and that further enhancement was not evident (Figure 2G, EPSP_kv4.2_ vs EPSC_kv4.2_, p>0.53) (Figure 2G). In spite, the effects of Kv4.2 inhibitions on the intrinsic excitability of GCs were evident. Kv4.2 Ab induced a significant increase in input resistance (219.5 ± 7.17 vs 253.97 ± 7.35, n = 67, p < 0.0001) and led to depolarization of the resting membrane potential (RMP) (−76.8 ± 0.79 vs −71.9 ± 0.93, n = 85, p < 0.0001) (Figure 2H, 2I). In contrast, AP half-width at rheobase current (0.75 ± 0.02 vs 0.74 ± 0.2, n = 16, p>0.7,) and the number of action potentials (AP) generated by +200 pA depolarizing current injections (200 pA: 7.76 ± 0.73 vs 8 ± 0.78, n = 16, p>0.5) showed no significant alteration due to Kv4.2 inhibitions, indicating that Kv4.2 had no discernible impact on GC firing properties (Figure 2J, 2K). Therefore, our findings imply that despite a substantial increase in input resistance by Kv4.2 inhibitions, this increase was not the predominant factor that induced amplification in EPSP amplitude, which contradicted to the well-known mechanism of K channel inhibition on EPSPs. Our data suggested that Kv4.2 channels exert profound influence on the AMPA-mediated currents to modulate synaptic response.

### Synaptic strength regulation by Kv4.2 channels correlates with its expression pattern in the hippocampus

To investigate the potential differential effects of Kv4.2 inhibitions across dendrites, we strategically placed the stimulation electrode to examine the effect of Kv4.2 inhibitions on EPSC amplitudes at different dendritic locations. In GCs, we observed no significant differences in the extent of EPSC enhancement induced by Kv4.2 Ab across different synaptic sites in GCs (Figure 3A, F (2,83) = [0.49], n = 6 for MML stimulation and n = 5 for IML stimulation, p = 0.61). However, CA3 PN exhibited distinct responses to Kv4.2 inhibitions depending on the input location. Stimulation in the stratum lucidum (SL) region showed no change in EPSC amplitude throughout the recording period with Kv4.2 Ab (Figure 3A, 1.00 ± 0.06, n = 5, p > 0.92). Conversely, Kv4.2 inhibition induced enhancement in EPSC amplitude when stimulating inputs in stratum radiatum (SR) (Figure 3B, 1.44 ± 0.12, n = 5, p < 0.05; EPSC_SL_ vs EPSC_SR_, p < 0.01).

**Figure 3.**
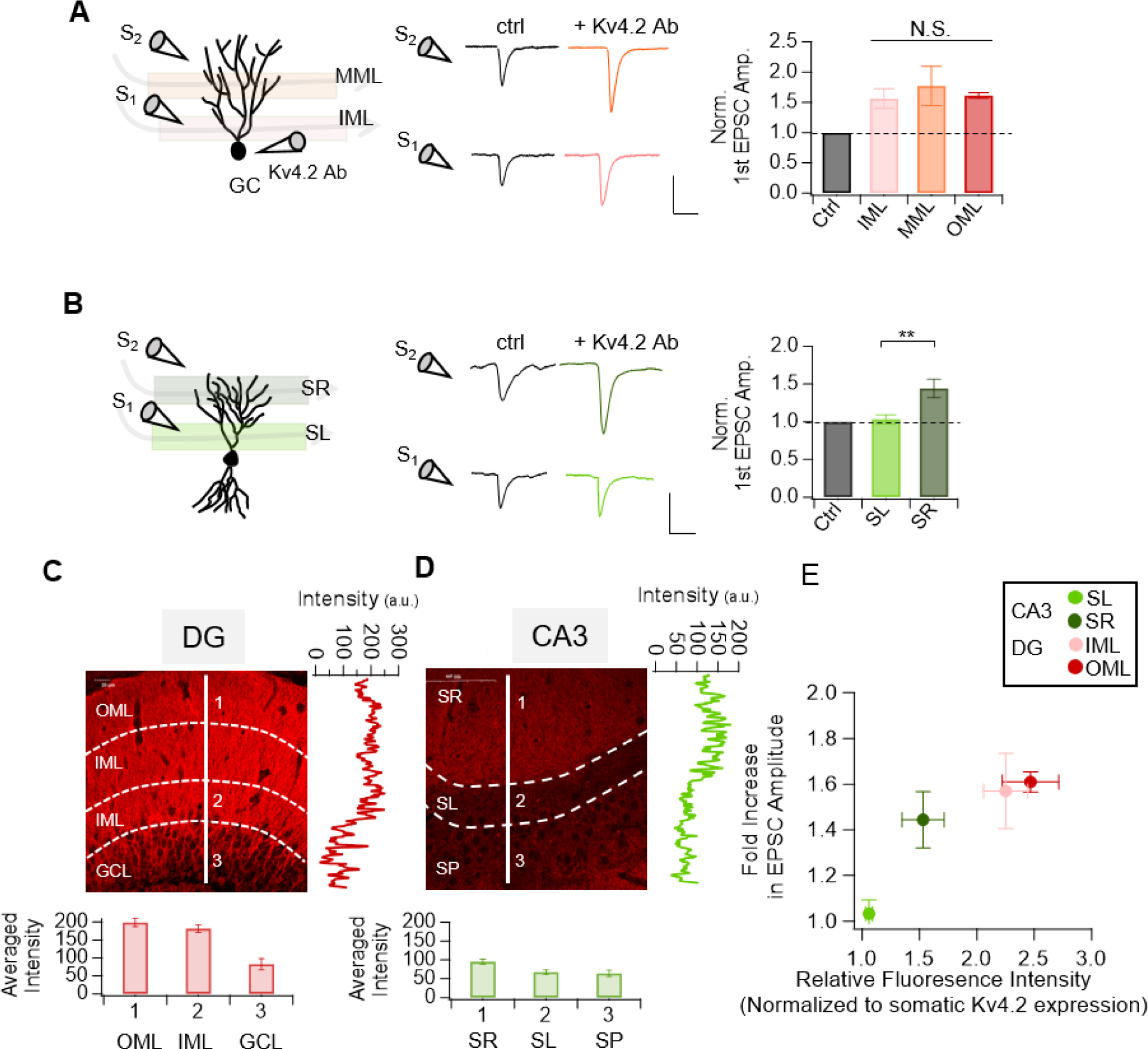
Magnitude of EPSC enhancement correlates with its expression pattern in the hippocampus. **A.** (Left) Recording configuration depicting the dialysis of either Kv4.2 Ab through recording pipette, with the stimulating electrode placed in IML (inner molecular layer) or MML (middle molecular layer) region of the dentate gyrus. (Middle) Representative average EPSC traces at 3-5 minutes (ctrl) and at 15-30 minutes (+Kv4.2 Ab) after patch break-in by stimulating MML region (Top) and IML region (Bottom) of DG. Scale bars, 100pA and 20ms. (Right) Bar graph illustrating the comparable enhancement of EPSC amplitude resulting from the inclusion of Kv4.2 Ab in the intracellular pipette in different synapses in GCs. **B.** (Left) Recording configuration depicting the dialysis of either Kv4.2 Ab through recording pipette, with the stimulating electrode placed in SL (stratum lucidum) or SR (stratum radiatum) region of the CA3. (Middle) Representative average EPSC traces at 3-5 minutes (ctrl) and at 15-30 minutes (+Kv4.2 Ab) after patch break-in by stimulating SR regions (Top) and SL regions (Bottom) of CA3 PN. Scale bars, 100pA and 20ms. (Right) Bar graph illustrating the enhancement of EPSC amplitude resulting from the inclusion of Kv4.2 Ab in the intracellular pipette in different synapses in CA3. **C.** Top: (Left) Representative fluorescence immunostaining images that show Kv4.2 expressions in DG of 7-week old mice. (Right) The representative intensity profile in the indicated white solid line in the image in the left. Bottom: Average fluorescence intensity in the marked number. **D.** Top: (Left) Representative fluorescence immunostaining images that show Kv4.2 expressions in CA3 of 7-week old mice. (Right) The representative intensity profile in the indicated white solid line in the image in the left. Bottom: Average fluorescence intensity in the marked number. **E.** Relative fluorescence intensity is plotted against EPSC enhancement induced by Kv4.2 inhibitions in different cell types measured. All data are represented as mean ± SEM. Statistical significance was evaluated by paired t tests, two-sample t-test, one-way ANOVA. **p<0.01, ****p<0.0001, n.s= not significant

Subsequently, we investigated whether different effects of Kv4.2 inhibitions on different synapses are related to the expression patterns of Kv4.2 in dendrites by conducting immunohistochemical analysis in CA3 and DG regions of the hippocampus. While both regions exhibited rare expression of Kv4.2 in somatic regions, robust Kv4.2 expressions was observed along the dendritic regions of GCs (Figure 3C), but not in CA3 (Figure 3D). In CA3 region, Kv4.2 expressions were relatively low in stratum lucidum (SL) but substantially higher in the stratum radiatum (SR) region (Figure 3D).

When the relative fluorescence intensity of Kv4.2 expressions in different dendritic layers was plotted against their magnitude of enhancement in EPSC amplitude, it was clear that Kv4.2 channel expression correlated with the extent of EPSC enhancement (Figure 3E). Dendrites in the SR, IML and OML, which exhibited relatively similar level of Kv4.2 expressions, showed comparable enhancement of EPSC amplitudes upon Kv4.2 inhibitions. In contrast, dendrites in SL in CA3, where Kv4.2 expressions was minimal, showed no potentiation in synaptic strength by Kv4.2 inhibitions. Therefore, our findings demonstrated Kv4.2 expressions correlated with synaptic strength regulation by Kv4.2.

### Kv4.2 inhibition-induced synaptic potentiation requires the R-type calcium channel in the dentate granule cells

To examine whether Kv4.2-dependent EPSC enhancement in LPP-GC synapses is Ca^2+^ dependent, we first introduced fast Ca^2+^ chelator 10 mM BAPTA with Kv4.2 Ab in the intracellular solution. Co-application of BAPTA and Kv4.2 Ab abolished EPSC enhancement (Figure 4A, 1.12 ± 0.06, n = 7, p > 0.12) without affecting RMP depolarization (Figure 4B, RMP_0.1EGTA_ vs RMP_10mMBAPTA,_ p > 0.05). These results suggested that Kv4.2 inhibitions directly induced RMP depolarization while it induced EPSC enhancement via Ca^2+^ signaling (Figure 4B).

**Figure 4.**
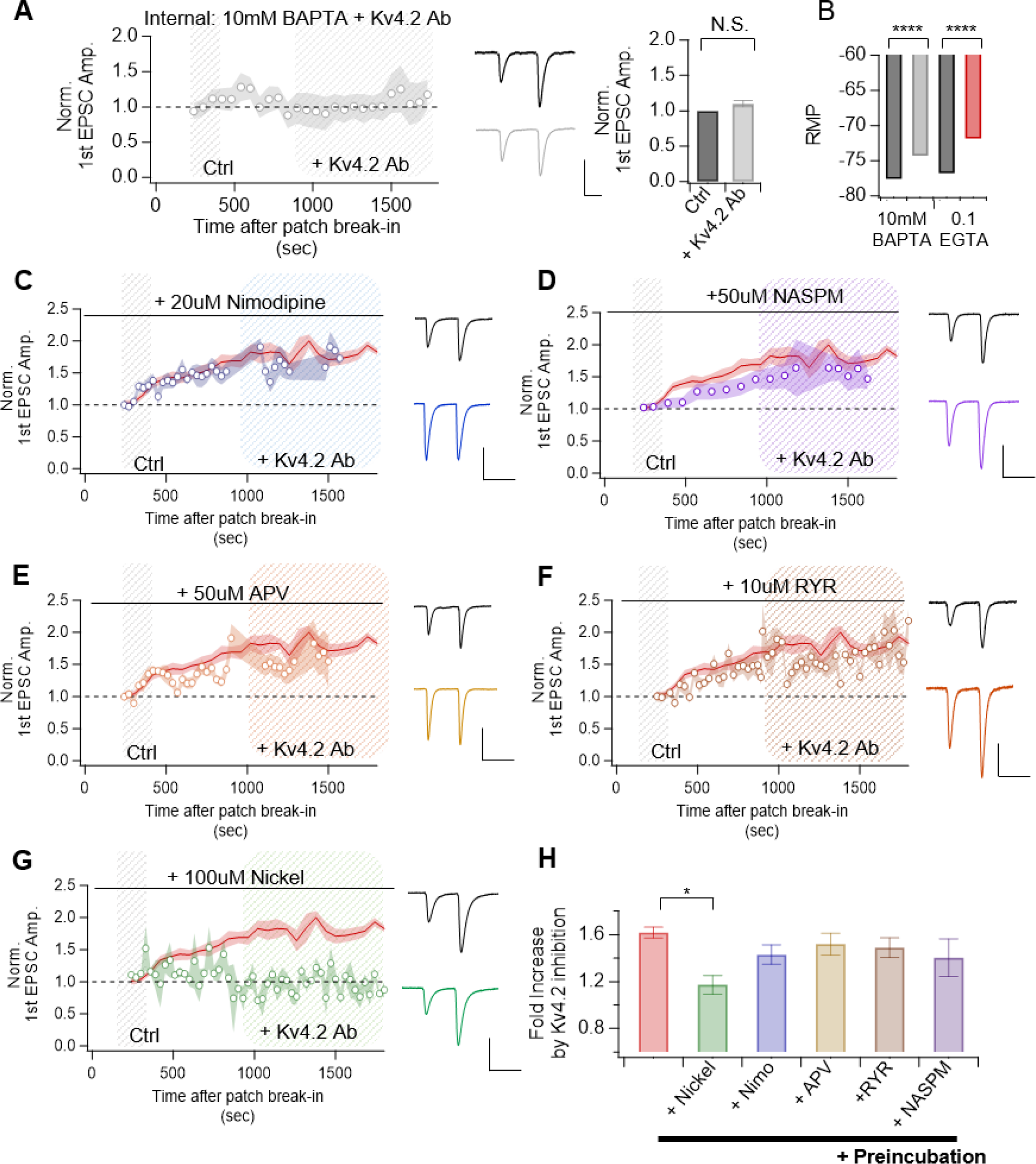
EPSC enhancement induced by Kv4.2 channel inhibition requires Ca2+ influx through R-type voltage-gated calcium channel in dentate granule cells. **A.** Time course summary plot (left) representative traces (middle) and summary bar graph (right) showing that EPSC enhancement is prevented by including 10mM BAPTA in intracellular solution. Scale bar, 50pA and 20ms. **B.** Changes in RMP induced by Kv4.2 inhibitions in the presence of 0.1 EGTA or 10mM BAPTA in intracellular solution **C-G**. Time course summary plot (left) and representative traces (right) showing effect of different calcium channels blockers on Kv4.2 Ab-induced EPSC enhancement. Kv4.2 inhibition induced EPSC enhancement is not abolished by the preincubating the slices (bath application for 20minutes before recording) with L-type calcium channel blocker Nimodipine (20uM) (B), CP-AMPAR blocker NASPM (50uM) (C), NMDAR blocker APV (50uM) (D) and Ryanodine receptor blocker RYR (10uM). Preapplication of R-type calcium channel (100uM bath application for 20 min) blocked Kv4.2-dependent enhancement in EPSC amplitude. Scale bar, 100pA and 50ms. **H**. Summary bar graph demonstrating that R-type calcium channel act as significant calcium source for Kv4.2 inhibition induced synaptic potentiation in GCs. Scale bar, 100pA and 50ms. All data are represented as mean ± SEM. Statistical significance was evaluated by paired t tests. ****p<0.0001, n.s= not significant

To identify the potential Ca^2+^ sources involved, we pretreated the slices with specific inhibitors for various postsynaptic Ca^2+^ sources, including NMDA receptors, voltage-gated Ca2+ channels (VGCCs), calcium-permeable AMPARs (CP-AMPARs), and ryanodine receptors (RYR) before assessing changes in EPSC amplitude by Kv4.2 Ab. Preincubation with APV (50 μM, NMDAR), nimodipine (10 μM, LTCC), NASPM (10 μM, CP-AMPAR) or ryanodine (10 μM, RYR) did not block Kv4.2 Ab-induced EPSC enhancement (Figure 4C-4F,4H)

However, the blockade of the R-type calcium channel (RTCC) with Ni^2+^ (100 μM) significantly inhibited the increase in EPSC by Kv4.2 Ab (Figure 4G, 1.12 ± 0.07, n = 16, p>0.05). Interestingly, low concentration of Ni^2+^ (30 μM), which is known to inhibit T-type calcium channels (TTCCs), did not prevent the potentiating action of Kv4.2 Ab (Figure 4 - supplement 1). Collectively, our findings provide evidence that potentiating action of Kv4.2 inhibitions on synaptic strength is dependent on RTCC-mediated Ca^2+^ signaling.

### Kv4.2 inhibition-induced synaptic potentiation is dependent on various signaling pathways

Numerous studies have shown that density of AMPARs can be regulated by various signaling pathways (Diering and Huganir., 2014). We next attempted to determine what kinases are involved to induce Kv4.2-dependent synaptic strengthening in PP-GCs. In the presence of blockers targeting phospholipase C (PLC) (10 μM, U73122) and Ca2+/calmodulin-dependent kinase (CaMKII) activities (5 μM, KN93), substantial EPSC enhancement by Kv4.2 inhibitions was still evident (Figure 5A-B, 5G). Furthermore, inclusion of calmodulin inhibitory peptide (CaM-IP) in intracellular solution failed to prevent Kv4.2 Ab-dependent EPSC enhancement, suggesting that synaptic strengthening induced by Kv4.2 Ab is independent from PLC and CaMKII signaling (Figure 5C, 5G).

**Figure 5.**
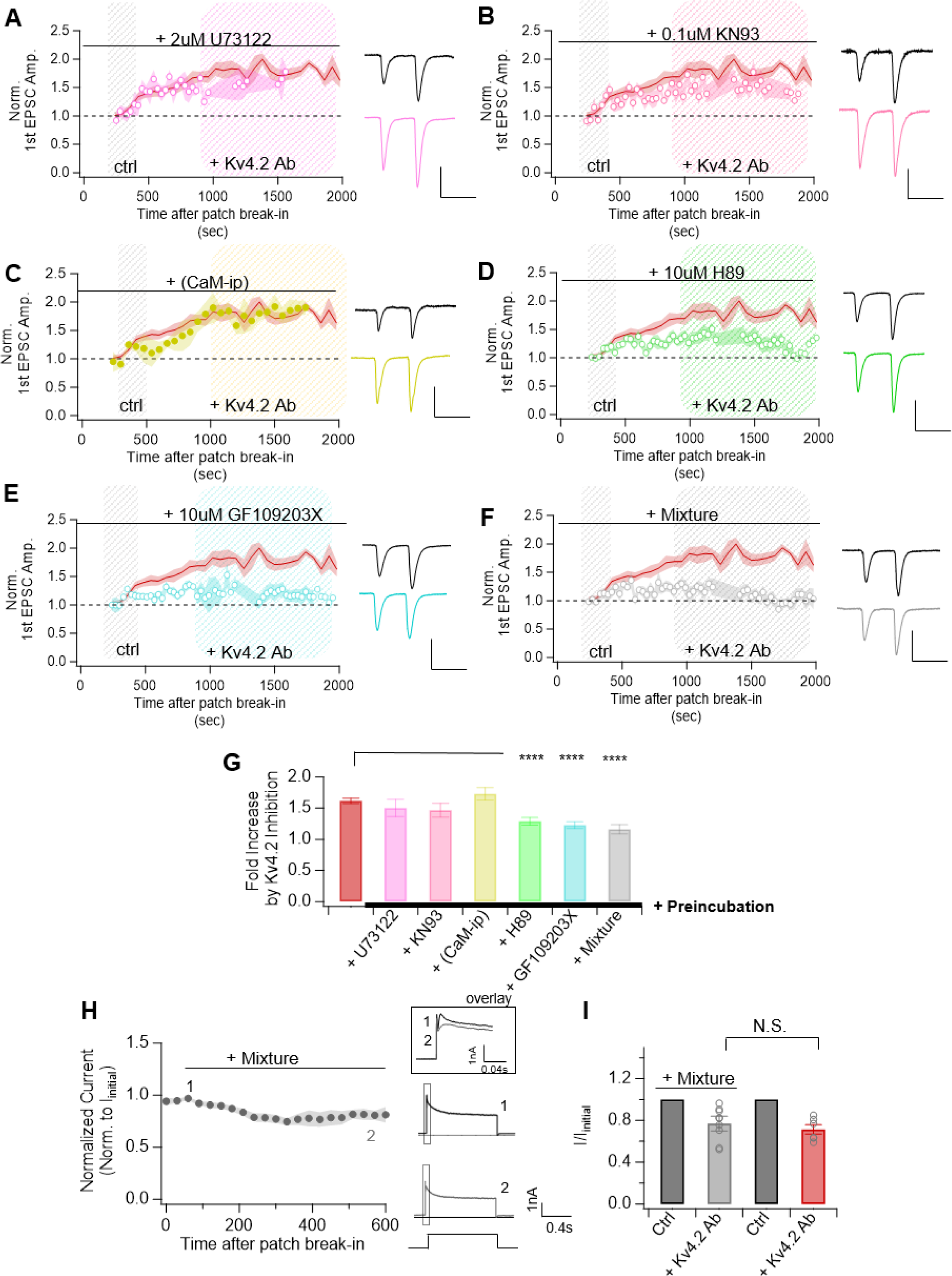
EPSC enhancement induced by Kv4.2 channel inhibition requires Ca^2+^ influx through R-type voltage-gated calcium channel in dentate granule cells. **A-B**. Time course summary plot (left) and representative traces (right) showing effect of different signaling pathway blockers on Kv4.2 Ab-induced EPSC enhancement. Kv4.2 inhibition induced EPSC enhancement is not abolished by the preincubating the slices (bath application for 20minutes before recording) with PLC inhibitor U73122 (2uM) (A) and CaMKII blocker KN93 (0.1uM) (B) in dentate granule cells of the hippocampus. Scale bar, 100pA, 50ms. **C-F**. Time course summary plot (left) and representative traces showing effect of different signaling pathway blockers on Kv4.2 Ab-induced EPSC enhancement. Kv4.2 inhibition induced EPSC enhancement is partially blocked by preincubating the slices (bath application for 20minutes before recording) with PKA inhibitor H89 (10uM) (C), PKC inhibitor GF109203X (10uM) (D) and calmodulin inhibitor calmidazolium (10uM) (E) in dentate granule cells of the hippocampus. In the presence of mixture (H89, GF109203X, and CMZ), Kv4.2-inhibition induced EPSC enhancement was completely blocked. Scale bar, 100pA, 50ms. **G**. Summary bar graph demonstrating that multiple signaling pathways are required for Kv4.2 inhibition induced synaptic potentiation in GCs. **H_a_.** The peak current amplitude was normalized to their respective initial value following patch break-in. These normalized amplitudes were plotted against the duration of the experiment. **H_b_**. Changes in transient current obtained at 10 min after patch break-in with or without the presence of mixture were normalized to their respective initial current amplitude and were plotted as bars with SEM. All data are represented as mean ± SEM. Statistical significance was evaluated by paired t tests. ****p<0.0001, n.s= not significant

Notably, blocking activity of protein kinase A (PKA) and protein kinase C (PKC) with H89 (10 μM) or GF109203X (10 μM) respectively, resulted in a significant yet partial reduction in EPSC enhancement (Figure 5C-E, 5G, H89: 1.28 ± 0.10, n = 21, p < 0.05; GF109203X: 1.23 ± 0.07, n = 20, p < 0.01). The combined preapplication of H89 and GF109203X entirely abolished the enhancement in EPSC amplitude induced by Kv4.2 Ab (Figure 5F-G, 1.15 ± 0.07, n = 9, p = 0.18). The possibility that these blockers affected A-type current was ruled out by confirming that Kv4.2 Ab-induced reduction of A-type current was comparable to the effects when Kv4.2 Ab were applied alone (Figure 5H, 0.77 ± 0.07, n = 9, p < 0.01; Transient_kv4.2_ vs Transient_Kv4.2 + mixture_, p>0.44). Overall, these findings suggest postsynaptic PKA and PKC activities are necessary for Kv4.2-dependent synaptic strengthening in GCs.

### Kv4.2-inhibition induced change in postsynaptic AMPAR in GCs

To further analyze the mechanism underlying the enhancement of AMPAR-mediated current induced by Kv4.2 Ab in GCs, we examined changes in the properties of spontaneous EPSC (sEPSC). Dialysis of Kv4.2 Ab resulted in a significant increase in sEPSC amplitude (Figure 6A_b_, 16.56 ± 0.46 vs 19.27 ± 0.61, n = 33, p < 0.0001), indicating an increase in the AMPAR density in postsynaptic site. Interestingly, sEPSC frequency was increased by Kv4.2 Ab in parallel (Figure 6Ac, 1.43 ± 0.13 vs 2.33 ± 0.26, n = 33, p > 0.0001).

**Figure 6.**
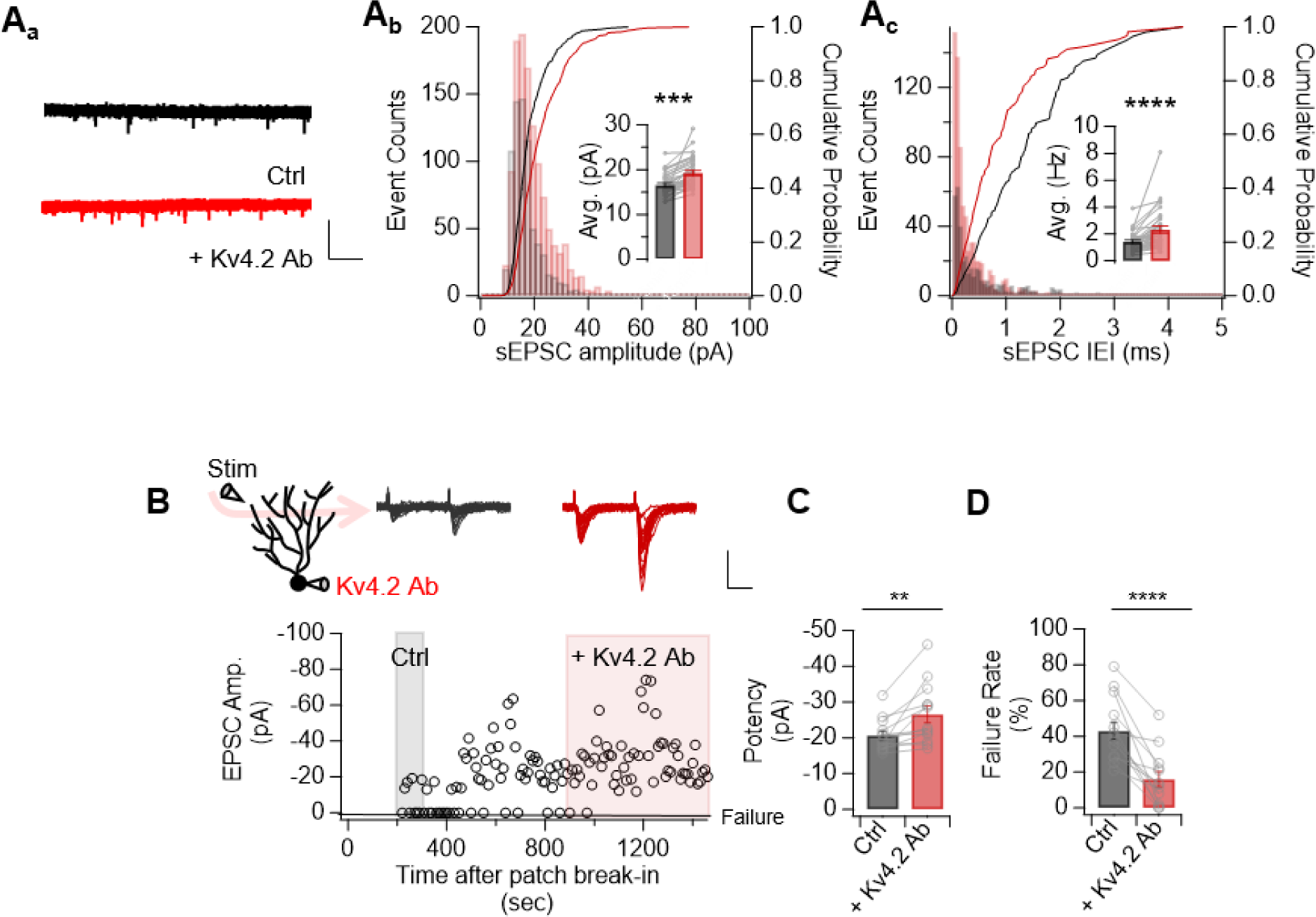
Kv4.2 inhibitions not only enhance AMPA density but also increase the number of synapses in dentate granule cells. **A_a_**. Representative spontaneous EPSC trace recorded at 5 minutes (ctrl) and at 30 minutes (+Kv.4 2Ab) after patch break-in with intracellular solution loaded with Kv4.2 Ab Scale bar, 50pA and 2s. **A_b_**. Histogram and cumulative plot showing changes induced in sEPSC amplitude in dentate granule cells by Kv4.2 Ab dialysis. Inset: bar graph showing enhanced sEPSC amplitude. **A_c_**. Histogram and cumulative plot showing changes induced in sEPSC frequency in dentate granule cells by Kv4.2 Ab dialysis. Inset: bar graph showing enhanced sEPSC frequency **B**. Representative experiment using minimal stimulation in OML; time course (bottom) and sample traces of control (ctrl, gray) and under the influence of Kv4.2 Ab (+Kv4.2 Ab, red) obtained from respective shaded area (top). Scale bar, 40pA, 20ms. **C-D**. Bar graph demonstrating that Kv4.2 inhibitions was associated significant increase in potency (C) and decrease failure rate (D). All data are represented as mean ± SEM. Statistical significance was evaluated by paired t tests. **p < 0.01, ****p<0.0001, n.s= not significant

Given that the NMDA-EPSC was intact following Kv4.2 inhibitions, involvement of presynaptic mechanism in Kv4.2 Ab-mediated synaptic strengthening could be ruled out. In spite that changes in sEPSC frequency are regarded as an indication of presynaptic modification, recent research proposed that the trafficking of AMPARs to NMDAR-containing silent synapses could influence sEPSC frequency by modifying the number of functional synapses at postsynaptic sites (Ying Yang et al., 2013). To test this possibility, we conducted minimal stimulation experiments to examine changes in eEPSC amplitude and failure rate by stimulating LEC synapses. Kv4.2 inhibitions led to an increase in potency (EPSC amplitude without failures) and a reduction in the failure rate (Figure 6B – 6D, potency: 1.42 ± 0.12, n = 16, p < 0.001; failure rate: 0.48 ± 0.05 vs 0.18 ± 0.05, n = 16, p < 0.0001). Therefore, enhanced sEPSC frequency and reduced failure rate by Kv4.2 inhibition suggest that AMPARs have been inserted into silent synapses of GCs. Overall, our results offer evidence that Kv4.2 inhibitions increase postsynaptic Ca^2+^ influx through RTCC to initiate PKA and PKC signaling events that ultimately leads to the postsynaptic insertion of AMPARs.

### Kv4.1 regulates dendritic excitability, while Kv4.2 regulates synaptic strength

Since Kv4.1 also regulates K^+^ conductance, we next attempted to examine whether Kv4.1 regulates synaptic strength. As previously reported (Kim et al., 2020), immunohistochemical analysis revealed a stronger expression of Kv4.1 in perisomatic regions of GCs, gradually decreasing along the proximo-distal axis of dendrites. Furthermore, expression of Kv4.1 in the dendritic regions was lower than the perisomatic region, with gradual decrease in the intensity along the proximo-distal axis of the dendrites (Figure 7A). The contrasting expression pattern of the Kv4.1 (Figure 7A) and Kv4.2 (Figure 3A) in the dendritic layers of hippocampal dentate gyrus suggested different roles of these two channels in synaptic transmission regulation.

**Figure 7.**
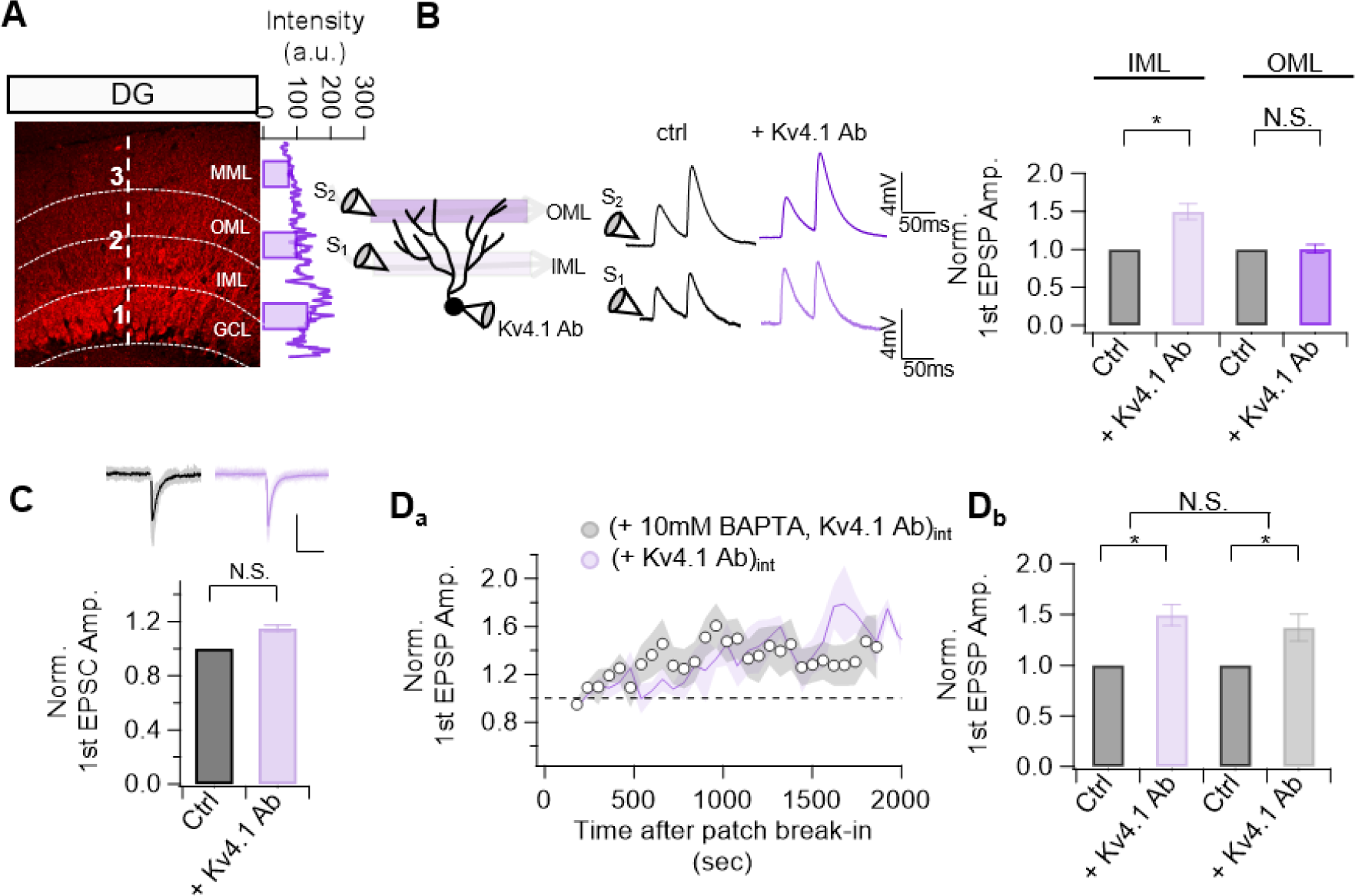
Synaptic strength regulation is specific to Kv4.2 channel. **A.** Top: (Left) Representative fluorescence immunostaining images that show Kv4.1 expression in DG of 7-week old mice. (Right) The representative intensity profile in the indicated line in the image in the left and the bar graph showing average fluorescence intensity in the marked number. **B**. Top: (Left) Recording configuration depicting the dialysis of either Kv4.2 Ab through recording pipette, with the stimulating electrode placed in OML or IML region of the dentate gyrus. (Middle) Representative average EPSP traces at control and at condition where Kv4.1 Ab reached its effect Scale bars, 4mV and 50ms. (Right) Bar graph illustrating the enhancement of EPSP amplitude resulting from the inclusion of Kv4.1 Ab in the intracellular pipette by placing stimulating electrode either in IML (light purple) or in OML (dark purple) **C** Top: Representative average EPSC traces at control and at condition where Kv4.1 Ab reached its effect Scale bars, 50pA and 50ms. Bottom: Bar graph showing changes in EPSC amplitude by Kv4.1 Ab **D_a_**. Time course of normalized EPSP amplitude in internal solution containing Kv4.2 with (represented by lines) or without 10mM BAPTA (represented by open circle). EPSP amplitudes were normalized to mean amplitude of the synaptic responses recorded at 3-5 minutes after patch break-in. Kv4.1 Ab induced increased EPSP amplitude even when 10mM BAPTA was included in intracellular solution. Data are binned into 30ms intervals, mean ± SEM. **D_b_.** Summary bar graph illustrating the enhancement of EPSP amplitude resulting from the inclusion of Kv4.2 Ab in the intracellular pipette is not different when 10mM BAPTA is included in intracellular solution. All data are represented as mean ± SEM. Statistical significance was evaluated by paired t tests. **p < 0.05, n.s= not significant

In order to assess the role of Kv4.1 on synaptic transmission, Kv4.1 Ab (∼ 1 μg/ml) was introduced into the intracellular solution and its effect on EPSP amplitude in LPP-GCs was examined. Unlike Kv4.2 inhibitions, there was no discernible change in EPSP amplitude was induced by Kv4.1 Ab when distal synapse in the OML regions were stimulated (Figure 7B, 1.01 ± 0.06, n = 9, p > 0.9). Interestingly, however, EPSPs evoked by stimulating proximal synapses in IML regions were significantly enhanced by Kv4.1 Ab, aligning with its subcellular expression pattern (Figure 7B, 1.49 ± 0.1, n = 6, p<0.01). This result further supported the functional link between K^+^ channel expression pattern and synaptic strength regulation

Interestingly, Kv4.1 inhibition induced no change in EPSC amplitude in proximal synapses of GC, indicating that Kv4.1 regulates dendritic excitability rather than synaptic strength (Figure 7C, 1.16 ± 0.03, n = 6, p > 0.13). This notion was further supported by minimal effect of 10 mM BAPTA on the Kv4.1 Ab-induced EPSP enhancement. Notably, co-application of BAPTA and Kv4.1 resulted in a substantial increase in EPSP amplitude, effect comparable to when Kv4.1 was applied alone (Figure 7D, 1.37 ±. 0.13, n = 7, p < 0.05; EPSP_Kv4.1_ vs EPSP_Kv4.1 + BAPTA_, p>0.8). Overall these data suggest that while Kv4.1 inhibition at proximal dendrite directly increases dendritic excitability to potentiate EPSPs in GCs, Kv4.2 inhibition increases synaptic strength via Ca^2+^-dependent regulation of AMPARs.

### Distinct mechanisms involved in Kv4.2 inhibition-induced synaptic strengthening in GCs

Long-term potentiation (LTP) of synaptic strength has been known to be attributed to increased density of AMPA receptors and downregulation of voltage-gated K^+^ channels that would synergistically produce potentiate synaptic responses. Therefore, we investigated the effect of Kv4.2 inhibitions on synaptic strengthening during LTP induction using theta burst stimulation (TBS). LTP experiments were initiated after 15 minutes of patch break-in to ensure that Kv4.2 Ab effect on AMPAR-mediated currents stabilized. After recording baseline EPSCs in LPP-GC synapses for about 5 minutes, 10 bouts of high frequency stimulation (HFS, 10 stimuli at 100 Hz) at 5 Hz was applied. EPSCs were subsequently measured for 15 minutes to observe the extent of synaptic potentiation induced by TBS. Even under conditions where Kv4.2 was inhibited, TBS still significantly increased EPSC amplitude (Figure 8A, 1.74 ± 0.15, p < 0.05). This enhancement in EPSC amplitude following LTP induction was comparable to that of the control groups, which did not include Kv4.2 Ab (Figure 8C, EPSC_+Kv4.2 Ab_ vs EPSC_-kv4.2Ab_: 1.85 ± 0.33 vs 1.74 ± 0.15, n = 4, p>0.6). Therefore, inhibition of Kv4.2 did not occlude further synaptic strengthening induced by TBS, indicating that the regulation of AMPAR by Kv4.2 differs from that of LTP.

**Figure 8.**
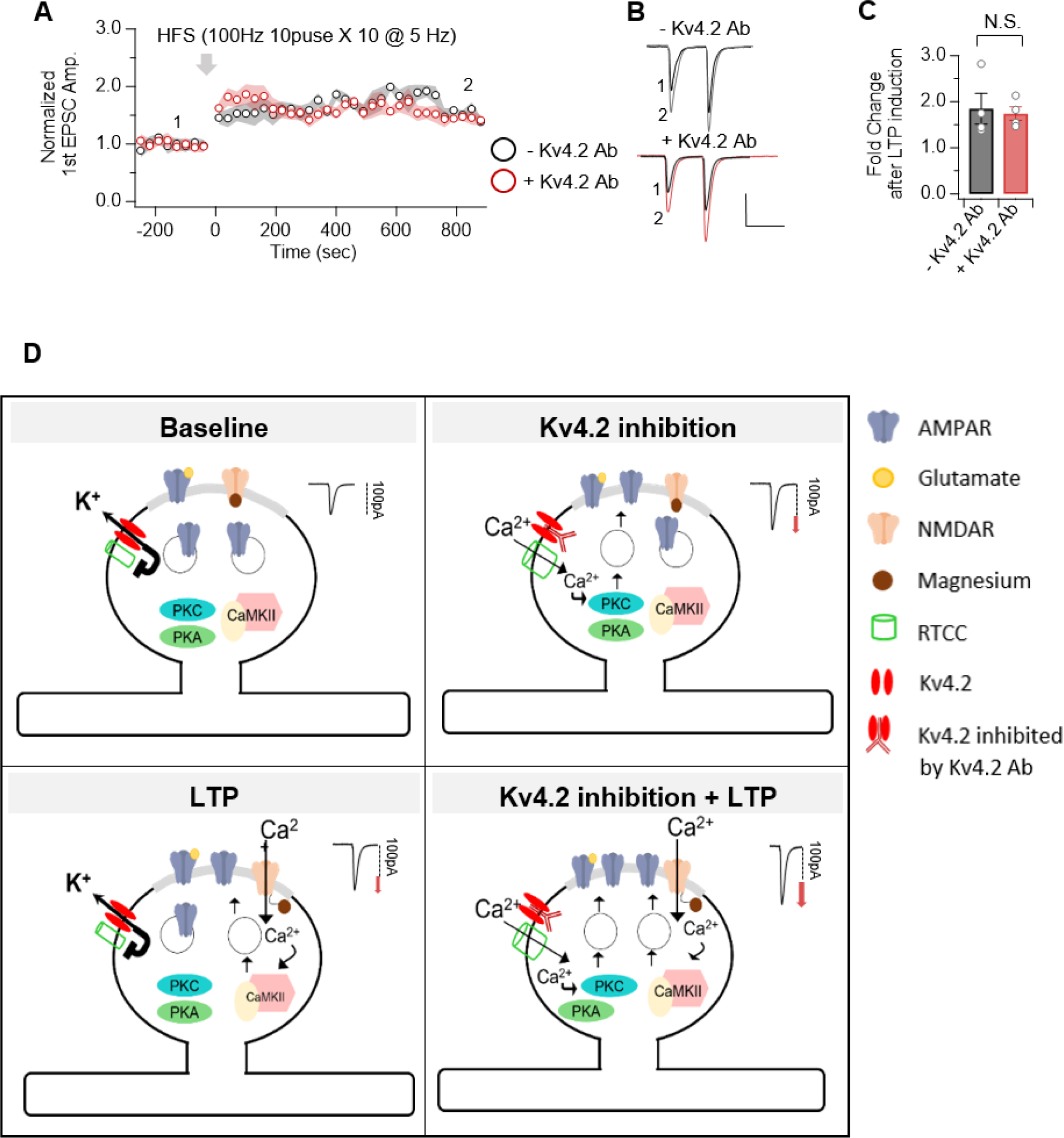
Kv4.2 channels do not occlude expression of LTP. **A.** Time course of normalized EPSC amplitude in internal solution containing Kv4.2 with (red) or without Kv4.1 Ab (black). Arrow indicates time point where HFS stimulation was given. HFS consisted of 10 bouts given at 5 Hz and each bout consisted of 100 Hz 10 pulse stimulation. Data are binned into 30ms intervals, mean ± SEM. **B.** Representative traces showing EPSC amplitude after LTP induction using internal solution with or without Kv4.2 Ab. **C.** Summary bar graph illustrating the enhancement of EPSC amplitude resulting from LTP induction is not different between the two groups. **D.** Summary illustration demonstrating the effect on Kv4.2 in synaptic transmission. During baseline synaptic transmission, Kv4.2 channels would suppress the activation of RTCC to limit the amount of Ca2+ influx the regulate appropriate level of AMPA density. Conversely, when Kv4.2 is inhibited, Ca2+ influx through RTCC would increase to activate PKA and PKC signaling molecules to enhance synaptic strength. The feedback mechanisms by Kv4.2-RTCC to control AMPA density is distinct from LTP because Kv4.2 inhibitions do not exclude the expression of further LTP, which necessitate the activation of NMDAR and CaMKII signaling pathways. All data are represented as mean ± SEM. Statistical significance was evaluated by paired t tests or two-sample t-test. **p < 0.05, n.s= not significant

## Discussion

In summary, as demonstrated in Figure 8D, our findings underscore that Kv4.2 channels regulate AMPAR-mediated currents by modulating RTCC-dependent Ca^2+^ signaling. Notably, Kv4.2 channel inhibition did not occlude the expression of LTP, suggesting that Kv4.2 Ab-induced synaptic strengthening operates through mechanisms distinct from EPSC enhancement observed during LTP induction. Consistent with these observations, RTCC and PKA/PKC emerged as primary regulators of Kv4.2-dependent synaptic strengthening, a process different from LTP induction predominantly governed by NMDAR and CaMKII (Hell et al., 2023). Also, Kv4.2 exhibited input-specific regulation depending on their expression pattern in the hippocampus. Overall, our findings proposed that Kv4.2 channels play essential role in maintaining baseline AMPA density through regulatory mechanism distinct from that involved in LTP.

### Kv4.2 channels affect AMPA receptor mediated current to shape synaptic response

Considerable evidence has established the role of Kv4.2-mediated A-type K^+^ current in dendritic signaling and synaptic plasticity (Hoffman et al., 1997; Kim et al., 2012, 2019, Kim et al., 2005; Jung et al., 2009; Simkim et al., 2015; Rathour et al., 2016; Carrasquillo et al., 2012; Oule et al., 2021; Hoffman et al., 1997). Inhibition of Kv4.2 mediated outward currents in dendrites have been regarded to amplify synaptic depolarization, but we could not find its amplification effect. On the contrary, our studies provide evidence that Kv4.2 channels regulate AMPAR-mediated currents to enhance EPSP amplitude.

To note, there are several shortcomings in methods that were used to assess Kv4.2 contributions in previous studies. Firstly, the commonly used transient A-type blocker, including 4-Amnopyridne (4-AP), BaCl_2_ or phrixotoxins (PaTx), has nonspecific effect on other potassium (Kv) channels, potentially complicating the interpretation of the obtained results. Secondly, both genetic ablation of Kv4.2 and pharmacological blockade of Kv4.2 channels fail to distinguish the effects of presynaptic and postsynaptic Kv4.2 channels. Additionally, compensatory overexpression of other K^+^ channels cannot be ruled out in studies using genetic ablation models. (Kim et al, 2005, Nerbonne et al., 2008; Foehring., 2008; Andrasfalvy et al., 2008). Altogether, it is difficult to interpret that those results are solely attributable to the blockade of dendritic Kv4.2 channels.

To circumvent these problems and identify the role of dendritic Kv4.2 channels in the synaptic responses of the cell, we recorded the synaptic responses with the patch clamp technique by including specific antibodies to Kv4.2 in the intracellular solution. Intracellular dialysis of Kv4.2 Ab inhibited transient current without affecting sustained current in GCs (Figure 1). By observing equivalent increase in EPSP and EPSC amplitude, we were able to conclude that Kv4.2 channels act as a powerful regulator of AMPA currents rather than dendritic excitability in hippocampal GCs (Figure 1, 2). It needs to be investigated in future studies whether other K^+^ channels known as dendritic channels can regulate AMPA currents.

### Regulation of RTCC by Kv4.2 inhibitions in dendritic spine

In our study, we have demonstrated that intracellular BAPTA and Ni^2+^ completely blocked potentiating effect of Kv4.2 Ab, indicating that Kv4.2-dependent regulation of AMPA currents is Ca^2+^ dependent. This finding strongly suggests that under basal conditions regulation of AMPA current by Kv4.2 channels is mediated by regulating calcium level via RTCCs. In support of this idea, it was previously shown that blockade of Kv4.2 channels led to an increase in spine calcium levels and mEPSC amplitude and these enhancements were prevented when R-type calcium channels were blocked using nickel (Ni^2+^) (Murphy et al., 2022).

Considering that L-type Ca^2+^ channel blocker did not prevent Kv4.2-dependent regulation of AMPA currents (Figure 4B), these suggested that Kv4.2-dependent regulation is not towards all voltage-gated Ca^2+^ channels, but specific to RTCCs. This specificity may be attributed to the fact that RTCC, not L-type calcium channel, is the major calcium source to action potential-evoked Ca^2+^ influx in dendritic spines in CA1 PN (Sabatini and Svobada 2000, Yasuda et al., 2003, Bloodgood and Sabatini et al., 2007). Also, the observed close proximity of Cav2.3 to Kv4.2 channels in dendritic spines of pyramidal neurons (Murphy et al., 2022) implicated that Kv4.2-mediated synaptic response regulation through RTCC is specialized function in dendritic spines. Given the considerable heterogeneity in the expression of VGCC classes across different cell types in dendritic spines, with cortical pyramidal neurons predominantly expressing L, P/Q, and T-type channels (Koester and Sakmann 2000), it is essential to explore the regulation of VGCCs by Kv4.2 channels in various cell types in the future studies.

In addition to VGCCs, NMDARs are potential calcium source at subthreshold potentials that can also be regulated by K^+^ channels (Sobczyk et al., 2005; Higley and Sabatini., 2008). For example, small-conductance calcium-activated channels (SK) in CA1 (Ngo Anh, 2005) and large-conductance calcium-activated channels (BK) channels in GCs (Zhang et al., 2018) regulate NMDAR activity in the dendritic spines. In the context, Ca^2+^ influx through NMDARs activate SK or BK channels and their activation provide hyperpolarizing current to inhibit NMDAR activation. Conversely, the inhibition of SK or BK channels enhance NMDA-mediated currents. However, we did not find any evidence that Kv4.2 inhibitions increase NMDARs-mediated currents. This may be due to higher activation threshold for NMDAR opening (Scheuss et al., 2009) than that of RTCC (Wormuth et al., 2016), suggesting that influence by Kv4.2 inhibitions at subthreshold potentials may be limited RTCC.

### RTCC-mediated regulation is distinct from NMDAR-dependent regulation for AMPA receptors

Our data revealed that AMPAR enhancement induced by Kv4.2 inhibition operates through distinct and nonoverlapping mechanisms compared to that associated with LTP induction (Figure 8). LTP in PP-GC synapses is known to regulated by NMDAR-(Kim et al., 2018) or LTCC-mediated (Lopez Rojas et al 2015, Kim et al., 2022) calcium influx, subsequently initiating CaMKII to increase AMPAR density in synaptic sites. However, Kv4.2-mediated regulation of synaptic strength primarily relied on RTCC-mediated calcium influx and signaling molecules including PKA and PKC

Furthermore, prior investigations have shown that inhibiting RTCC-mediated Ca^2+^ signaling reduced I_A_ current by decreasing the surface expression level of Kv4.2 in a KChIP-dependent manner (Wang et al, 2014, Murphy et al., 2022). These studies suggested that RTCC-mediated Ca^2+^ influx promotes the functional expression of Kv4.2 in dendrites (Murphy et al., 2022). Considering that our data provides evidence that RTCC-mediated Ca^2+^ influx contributes to enhancement in AMPAR-mediated current, RTCC-mediated Ca^2+^ signaling serves to regulate both AMPA density and Kv4.2 surface expression. These suggest that Ca^2+^ influx through RTCC enhances AMPA density while simultaneously increasing Kv4.2 surface expression, which may act as a mechanism to suppress further increments in AMPA density. Therefore, Kv4.2-RTCC may establish a negative feedback loop to maintain an appropriate level of synaptic Ca^2+^ and AMPARs under basal conditions.

This regulation of AMPAR and Kv4.2 by RTCC-mediated calcium influx differs from that by NMDAR during LTP. It is a widely established that NMDAR-mediated Ca2+influx leads to trafficking of AMPARs to surface membrane while it induces the internalization of Kv4.2, working synergistically to enhance EPSP amplitude (Kim and Hoffman., 2007, Jung and Hoffman., 2009). Therefore, while robust increase in AMPAR density in association with Kv4.2 inhibitions is induced by NMDAR-mediated calcium influx during LTP, AMPAR remains limited in the physiological range under baseline synaptic transmission by Kv4.2-RTCC negative feedback mechanism. In conclusion, given the broad expression of Kv4.2 in the brain and potential pathological impacts of disruptions in Kv4.2 function, our research offers a comprehensive framework for understanding the physiological and pathological roles of Kv4.2-dependent regulation of synaptic transmission.

---

**Figure 4-Supplement 1.**
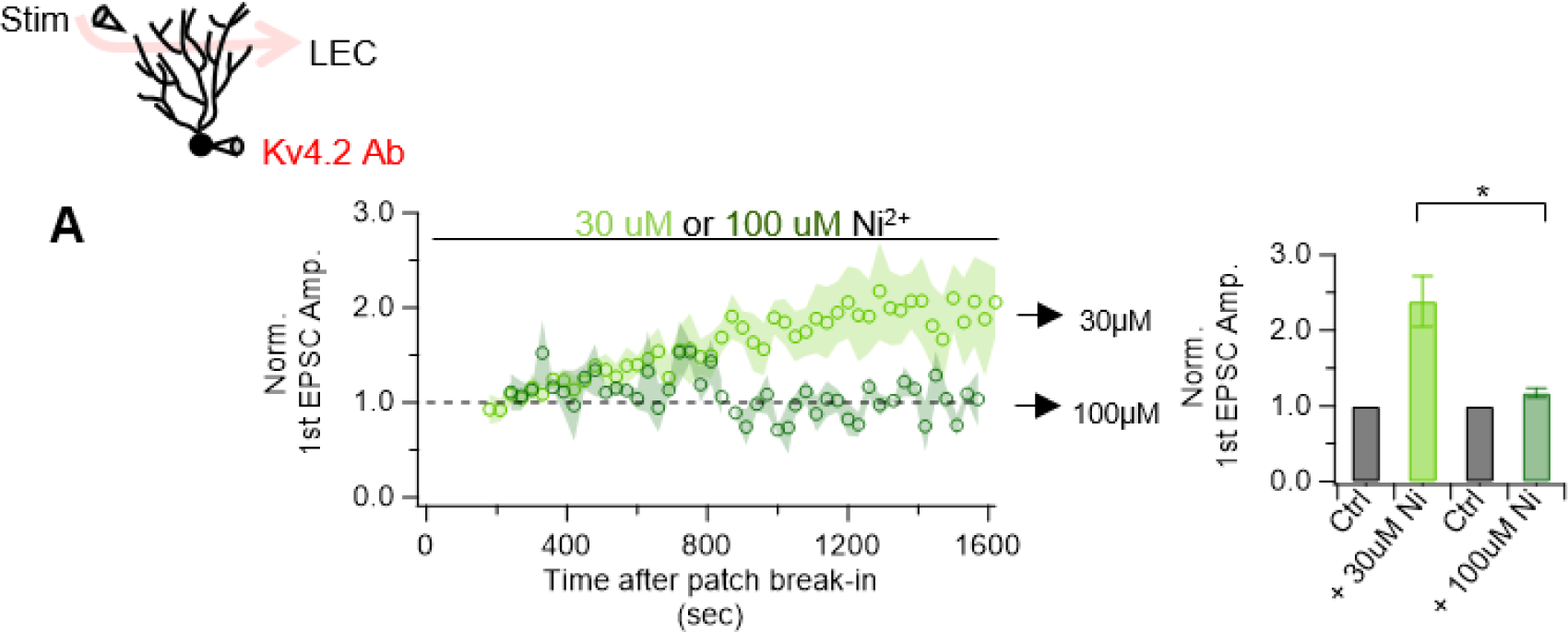
EPSC enhancement induced by Kv4.2 channel inhibition is inhibited by 100uM Nickel (Ni^2+^) **A.** Time course summary plot (left) and bar graph (left) showing that EPSC enhancement is prevented by 100uM Ni^2+^, not 30uM Ni^2+^. All data are represented as mean ± SEM. Statistical significance was evaluated by paired t tests. **p < 0.05, n.s= not significant

## References

1. Alfaro-Ruiz, R., Aguado, C., Martin-Belmonte, A., Moreno-Martinez, A. E., & Lujan, R. (2019). Expression, Cellular and Subcellular Localisation of Kv4.2 and Kv4.3 Channels in the Rodent Hippocampus. Int J Mol Sci, 20(2). doi:10.3390/ijms20020246

2. Andrásfalvy, B., Makara, J., Johnston, D., & Magee, J. (2008). Altered synaptic and non-synaptic properties of CA1 pyramidal neurons in Kv4. 2 knockout mice. The Journal of physiology, 586(16), 3881–3892.

3. Barcomb, K., Hell, J. W., Benke, T. A., & Bayer, K. U. (2016). The CaMKII/GluN2B protein interaction maintains synaptic strength. Journal of Biological Chemistry, 291(31), 16082–16089.

4. Bliss, T. V., & Collingridge, G. L. (2013). Expression of NMDA receptor-dependent LTP in the hippocampus: bridging the divide. Molecular brain, 6(1), 1–14.

5. Bloodgood, B. L., & Sabatini, B. L. (2008). Regulation of synaptic signalling by postsynaptic, non-glutamate receptor ion channels. The Journal of physiology, 586(6), 1475–1480.

6. Bredt, D. S., & Nicoll, R. A. (2003). AMPA receptor trafficking at excitatory synapses. Neuron, 40(2), 361–379.

7. Cai, X., Liang, C. W., Muralidharan, S., Kao, J. P., Tang, C.-M., & Thompson, S. M. (2004). Unique roles of SK and Kv4. 2 potassium channels in dendritic integration. Neuron, 44(2), 351–364. Retrieved from https://www.sciencedirect.com/science/article/pii/S089662730400635X?via%3Dihub

8. Carrasquillo, Y., Burkhalter, A., & Nerbonne, J. M. (2012). A-type K+ channels encoded by Kv4. 2, Kv4. 3 and Kv1. 4 differentially regulate intrinsic excitability of cortical pyramidal neurons. The Journal of physiology, 590(16), 3877–3890.

9. Chen, X., Yuan, L. L., Zhao, C., Birnbaum, S. G., Frick, A., Jung, W. E., Johnston, D. (2006). Deletion of Kv4.2 gene eliminates dendritic A-type K+ current and enhances induction of long-term potentiation in hippocampal CA1 pyramidal neurons. J Neurosci, 26(47), 12143–12151. doi:10.1523/JNEUROSCI.2667-06.2006

10. Chen, X., Yuan, L.-L., Zhao, C., Birnbaum, S. G., Frick, A., Jung, W. E., Johnston, D. (2006). Deletion of Kv4. 2 gene eliminates dendritic A-type K+ current and enhances induction of long-term potentiation in hippocampal CA1 pyramidal neurons. Journal of Neuroscience, 26(47), 12143–12151.

11. Diaz-Alonso, J., & Nicoll, R. A. (2021). AMPA receptor trafficking and LTP: Carboxy-termini, amino-termini and TARPs. Neuropharmacology, 197, 108710. doi:10.1016/j.neuropharm.2021.108710

12. Diering, G. H., & Huganir, R. L. (2018). The AMPA Receptor Code of Synaptic Plasticity. Neuron, 100(2), 314–329. doi:10.1016/j.neuron.2018.10.018

13. Faber, E. S., Delaney, A. J., & Sah, P. (2005). SK channels regulate excitatory synaptic transmission and plasticity in the lateral amygdala. Nat Neurosci, 8(5), 635–641. doi:10.1038/nn1450

14. Foehring, R. C. (2008). Who needs A current? Functional remodelling in the Kv4. 2−/− mouse. The Journal of physiology, 586(Pt 6), 1461.

15. Frick, A., Magee, J., & Johnston, D. (2004). LTP is accompanied by an enhanced local excitability of pyramidal neuron dendrites. Nat Neurosci, 7(2), 126–135. doi:10.1038/nn1178

16. Gómez, R., Maglio, L. E., Gonzalez-Hernandez, A. J., Rivero-Pérez, B., Bartolomé-Martín, D., & Giraldez, T. (2021). NMDA receptor–BK channel coupling regulates synaptic plasticity in the barrel cortex. Proceedings of the National Academy of Sciences, 118(35), e2107026118.

17. Hardingham, G. E., & Bading, H. (2010). Synaptic versus extrasynaptic NMDA receptor signalling: implications for neurodegenerative disorders. Nature reviews neuroscience, 11(10), 682–696.

18. Hell, J. W. (2023). Binding of CaMKII to the NMDA receptor is sufficient for long-term potentiation. Science Signaling, 16(808), eadk9224.

19. Herring, B. E., & Nicoll, R. A. (2016). Long-Term Potentiation: From CaMKII to AMPA Receptor Trafficking. Annu Rev Physiol, 78, 351–365. doi:10.1146/annurev-physiol-021014-071753

20. Higley, M. J., & Sabatini, B. L. (2008). Calcium signaling in dendrites and spines: practical and functional considerations. Neuron, 59(6), 902–913.

21. Hoffman, D. A., Magee, J. C., Colbert, C. M., & Johnston, D. (1997). K+ channel regulation of signal propagation in dendrites of hippocampal pyramidal neurons. nature, 387(6636), 869–875.

22. Johnston, D., Christie, B. R., Frick, A., Gray, R., Hoffman, D. A., Schexnayder, L. K., … Yuan, L.-L. (2003). Active dendrites, potassium channels and synaptic plasticity. Philosophical Transactions of the Royal Society of London. Series B: Biological Sciences, 358(1432), 667–674.

23. Jung, S. C., & Hoffman, D. A. (2009). Biphasic somatic A-type K channel downregulation mediates intrinsic plasticity in hippocampal CA1 pyramidal neurons. PloS one, 4(8), e6549. doi:10.1371/journal.pone.0006549

24. Jung, S.-C., & Hoffman, D. A. (2009). Biphasic somatic A-type K+ channel downregulation mediates intrinsic plasticity in hippocampal CA1 pyramidal neurons. PloS one, 4(8), e6549.

25. Kerloch, T., Clavreul, S., Goron, A., Abrous, D. N., & Pacary, E. (2019). Dentate granule neurons generated during perinatal life display distinct morphological features compared with later-born neurons in the mouse hippocampus. Cerebral cortex, 29(8), 3527–3539.

26. Kim, J., & Hoffman, D. A. (2008). Potassium channels: newly found players in synaptic plasticity. The Neuroscientist, 14(3), 276–286.

27. Kim, J., & Hoffman, D. A. (2008). Potassium channels: newly found players in synaptic plasticity. The Neuroscientist, 14(3), 276–286.

28. Kim, J., Wei, D. S., & Hoffman, D. A. (2005). Kv4 potassium channel subunits control action potential repolarization and frequency-dependent broadening in rat hippocampal CA1 pyramidal neurones. J Physiol, 569(Pt 1), 41–57. doi:10.1113/jphysiol.2005.095042

29. Kim, J. H., Sung-Cherl Jung, Ann M. Clemens, Ronald S. Petralia, and Dax A. Hoffman. (2007). Regulation of dendritic excitability by activity-dependent trafficking of the A-type K+ channel subunit Kv4.2 in hippocampal neurons. Neuron. doi: 10.1016/j.neuron.2007.05.026

30. Kim, K. R., Lee, S. Y., Yoon, S. H., Kim, Y., Jeong, H. J., Lee, S., … Ho, W. K. (2020). Kv4.1, a Key Ion Channel For Low Frequency Firing of Dentate Granule Cells, Is Crucial for Pattern Separation. J Neurosci, 40(11), 2200–2214. doi:10.1523/JNEUROSCI.1541-19.2020

31. Kim, S., Guzman, S. J., Hu, H., & Jonas, P. (2012). Active dendrites support efficient initiation of dendritic spikes in hippocampal CA3 pyramidal neurons. Nat Neurosci, 15(4), 600–606. doi:10.1038/nn.3060

32. Kim, S., Kim, Y., Lee, S. H., & Ho, W. K. (2018). Dendritic spikes in hippocampal granule cells are necessary for long-term potentiation at the perforant path synapse. elife, 7. doi:10.7554/eLife.35269

33. Koester, H. J., & Sakmann, B. (2000). Calcium dynamics associated with action potentials in single nerve terminals of pyramidal cells in layer 2/3 of the young rat neocortex. The Journal of physiology, 529(3), 625–646.

34. Kristensen, A. S., Jenkins, M. A., Banke, T. G., Schousboe, A., Makino, Y., Johnson, R. C., Traynelis, S. F. (2011). Mechanism of Ca2+/calmodulin-dependent kinase II regulation of AMPA receptor gating. Nature neuroscience, 14(6), 727–735.

35. Lei, Z., Deng, P., & Xu, Z. C. (2008). Regulation of Kv4. 2 channels by glutamate in cultured hippocampal neurons. Journal of neurochemistry, 106(1), 182–192.

36. Lin, M. T., Luján, R., Watanabe, M., Adelman, J. P., & Maylie, J. (2008). SK2 channel plasticity contributes to LTP at Schaffer collateral–CA1 synapses. Nature neuroscience, 11(2), 170–177.

37. Lisman, J., Yasuda, R., & Raghavachari, S. (2012). Mechanisms of CaMKII action in long-term potentiation. Nat Rev Neurosci, 13(3), 169–182. doi:10.1038/nrn3192

38. Lopez-Rojas, J., Heine, M., & Kreutz, M. R. (2016). Plasticity of intrinsic excitability in mature granule cells of the dentate gyrus. Scientific Reports, 6(1), 21615.

39. Losonczy, A., & Magee, J. C. (2006). Integrative properties of radial oblique dendrites in hippocampal CA1 pyramidal neurons. Neuron, 50(2), 291–307.

40. Luscher, C., & Malenka, R. C. (2012). NMDA receptor-dependent long-term potentiation and long-term depression (LTP/LTD). Cold Spring Harb Perspect Biol, 4(6). doi:10.1101/cshperspect.a005710

41. Migliore, M., Hoffman, D. A., Magee, J. C., & Johnston, D. (1999). Role of an A-type K+ conductance in the back-propagation of action potentials in the dendrites of hippocampal pyramidal neurons. Journal of computational neuroscience, 7, 5–15.

42. Murphy, J. G., Gutzmann, J. J., Lin, L., Hu, J., Petralia, R. S., Wang, Y.-X., & Hoffman, D. A. (2022). R-type voltage-gated Ca2+ channels mediate A-type K+ current regulation of synaptic input in hippocampal dendrites. Cell reports, 38(3).

43. Nerbonne, J. M., Gerber, B. R., Norris, A., & Burkhalter, A. (2008). Electrical remodelling maintains firing properties in cortical pyramidal neurons lacking KCND2-encoded A-type K+ currents. The Journal of physiology, 586(6), 1565–1579.

44. Neveu, D., & Zucker, R. S. (1996). Postsynaptic levels of [Ca2+] i needed to trigger LTD and LTP. Neuron, 16(3), 619–629.

45. Newpher, T. M., & Ehlers, M. D. (2008). Glutamate receptor dynamics in dendritic microdomains. Neuron, 58(4), 472–497.

46. Ngo-Anh, T. J., Bloodgood, B. L., Lin, M., Sabatini, B. L., Maylie, J., & Adelman, J. P. (2005). SK channels and NMDA receptors form a Ca2+-mediated feedback loop in dendritic spines. Nat Neurosci, 8(5), 642–649. doi:10.1038/nn1449

47. Opazo, P., Labrecque, S., Tigaret, C. M., Frouin, A., Wiseman, P. W., De Koninck, P., & Choquet, D. (2010). CaMKII triggers the diffusional trapping of surface AMPARs through phosphorylation of stargazin. Neuron, 67(2), 239–252. doi:10.1016/j.neuron.2010.06.007

48. Oulé, M., Atucha, E., Wells, T. M., Macharadze, T., Sauvage, M. M., Kreutz, M. R., & Lopez-Rojas, J. (2021). Dendritic Kv4. 2 potassium channels selectively mediate spatial pattern separation in the dentate gyrus. Iscience, 24(8).

49. Parajuli, L. K., Nakajima, C., Kulik, A., Matsui, K., Schneider, T., Shigemoto, R., & Fukazawa, Y. (2012). Quantitative regional and ultrastructural localization of the Cav2. 3 subunit of R-type calcium channel in mouse brain. Journal of Neuroscience, 32(39), 13555–13567.

50. Ramakers, G. M., & Storm, J. F. (2002). A postsynaptic transient K+ current modulated by arachidonic acid regulates synaptic integration and threshold for LTP induction in hippocampal pyramidal cells. Proceedings of the National Academy of Sciences, 99(15), 10144–10149.

51. Rathour, R., Malik, R., & Narayanan, R. (2016). Transient potassium channels augment degeneracy in hippocampal active dendritic spectral tuning. Sci Rep 6: 24678. In.

52. Rhodes, K. J., Carroll, K. I., Sung, M. A., Doliveira, L. C., Monaghan, M. M., Burke, S. L., … Cao, J. (2004). KChIPs and Kv4 α subunits as integral components of A-type potassium channels in mammalian brain. Journal of Neuroscience, 24(36), 7903–7915.

53. Rosenkranz, J. A., Frick, A., & Johnston, D. (2009). Kinase-dependent modification of dendritic excitability after long-term potentiation. J Physiol, 587(1), 115–125. doi:10.1113/jphysiol.2008.158816

54. Sabatini, B. L., & Svoboda, K. (2000). Analysis of calcium channels in single spines using optical fluctuation analysis. nature, 408(6812), 589–593.

55. Sanhueza, M., Fernandez-Villalobos, G., Stein, I. S., Kasumova, G., Zhang, P., Bayer, K. U., … Lisman, J. (2011). Role of the CaMKII/NMDA receptor complex in the maintenance of synaptic strength. J Neurosci, 31(25), 9170–9178. doi:10.1523/JNEUROSCI.1250-11.2011

56. Scheefhals, N., Westra, M., & MacGillavry, H. D. (2023). mGluR5 is transiently confined in perisynaptic nanodomains to shape synaptic function. Nature communications, 14(1), 244.

57. Simkin, D., Hattori, S., Ybarra, N., Musial, T. F., Buss, E. W., Richter, H., Disterhoft, J. F. (2015). Aging-Related Hyperexcitability in CA3 Pyramidal Neurons Is Mediated by Enhanced A-Type K+ Channel Function and Expression. J Neurosci, 35(38), 13206–13218. doi:10.1523/JNEUROSCI.0193-15.2015

58. Sobczyk, A., Scheuss, V., & Svoboda, K. (2005). NMDA receptor subunit-dependent [Ca2+] signaling in individual hippocampal dendritic spines. Journal of Neuroscience, 25(26), 6037–6046.

59. Tomita, S., Stein, V., Stocker, T. J., Nicoll, R. A., & Bredt, D. S. (2005). Bidirectional synaptic plasticity regulated by phosphorylation of stargazin-like TARPs. Neuron, 45(2), 269–277. doi:10.1016/j.neuron.2005.01.009

60. Varga, A. W., Anderson, A. E., Adams, J. P., Vogel, H., & Sweatt, J. D. (2000). Input-specific immunolocalization of differentially phosphorylated Kv4.2 in the mouse brain. Learn Mem, 7(5), 321–332. doi:10.1101/lm.35300

61. Wang, K., Lin, M. T., Adelman, J. P., & Maylie, J. (2014). Distinct Ca2+ sources in dendritic spines of hippocampal CA1 neurons couple to SK and Kv4 channels. Neuron, 81(2), 379–387. doi:10.1016/j.neuron.2013.11.004

62. Watanabe, S., Hoffman, D. A., Migliore, M., & Johnston, D. (2002). Dendritic K+ channels contribute to spike-timing dependent long-term potentiation in hippocampal pyramidal neurons. Proceedings of the National Academy of Sciences, 99(12), 8366–8371.

63. Watanabe, S., Hoffman, D. A., Migliore, M., & Johnston, D. (2002). Dendritic K+ channels contribute to spike-timing dependent long-term potentiation in hippocampal pyramidal neurons. Proceedings of the National Academy of Sciences, 99(12), 8366–8371.

64. Yang, K., & Dani, J. A. (2014). Dopamine D1 and D5 receptors modulate spike timing-dependent plasticity at medial perforant path to dentate granule cell synapses. J Neurosci, 34(48), 15888–15897. doi:10.1523/JNEUROSCI.2400-14.2014

65. Yang, S., Tang, C. M., & Yang, S. (2015). The Shaping of Two Distinct Dendritic Spikes by A-Type Voltage-Gated K(+) Channels. Front Cell Neurosci, 9, 469. doi:10.3389/fncel.2015.00469

66. Yang, Y. S., Kim, K. D., Eun, S. Y., & Jung, S. C. (2014). Roles of somatic A-type K(+) channels in the synaptic plasticity of hippocampal neurons. Neurosci Bull, 30(3), 505–514. doi:10.1007/s12264-013-1399-7

67. Yasuda, R., Hayashi, Y., & Hell, J. W. (2022). CaMKII: a central molecular organizer of synaptic plasticity, learning and memory. Nat Rev Neurosci, 23(11), 666–682. doi:10.1038/s41583-022-00624-2

68. Yasuda, R., Sabatini, B. L., & Svoboda, K. (2003). Plasticity of calcium channels in dendritic spines. Nature neuroscience, 6(9), 948–955.

69. Zhang, J., Guan, X., Li, Q., Meredith, A. L., Pan, H.-L., & Yan, J. (2018). Glutamate-activated BK channel complexes formed with NMDA receptors. Proceedings of the National Academy of Sciences, 115(38), E9006–E9014.

70. Zhao, C., Wang, L., Netoff, T., & Yuan, L. L. (2011). Dendritic mechanisms controlling the threshold and timing requirement of synaptic plasticity. Hippocampus, 21(3), 288–297.

